# Seventy-five Years of Systematic Biology: Looking Back, Moving Forward

**DOI:** 10.1101/2025.10.21.683502

**Authors:** Michael J. Landis, Michael J. Donoghue

## Abstract

What does “systematic biology” mean today, where has it been in the past, and where is it going? We explore these questions by considering five elements – collaboration, integration, discourse, infrastructure, and society – that we think have allowed systematic biology to adapt to change and sustain growth without losing its unique identity. In the spirit of celebrating the 75^th^ anniversary of our flagship journal, we generated a comprehensive dataset for all *Systematic Biology* and *Systematic Zoology* articles that we could locate (*N* = 5,150) and used bibliometric and textual analyses to illustrate ways in which our field has transformed over time. We offer our humble opinions on how our community can ensure that systematic biologists inherit an enlightening, dynamic, and enduring future.

---

We are truly grateful for this opportunity to reflect on the state of our discipline and of our flagship publication – *Systematic Zoology* turned *Systematic Biology* – as it enters its 75th year. It has been great fun for the two of us to chat through the current state of systematic biology, to look back at how it has changed over these past seven decades, and to think forward to what it may become. It is only on rare occasions like this one that we can step back and ask the sorts of existential (and perhaps indulgent) questions normally enjoyed in close company: What is systematic biology anyway? Is systematics in a stable phase of normal science, productive but perhaps stuck in a rut? Or is it instead undergoing, or about to undergo, a radical shift? And, what would any such answers mean for the health and the future of our discipline?

More than 35 years have passed since David Hull’s monumental *Science as a Process* (Hull, 1988) – his Thomas Kuhn-inspired analysis of the development of systematic biology – and a whole lot has happened since then. We began our musings in this Hullian spirit, though with ambitions greatly tempered by our own limited knowledge, and, of course, having been commissioned to write only a few pages on the topic. To help illustrate our reflections, we compiled a dataset containing all articles we could locate from Volumes 1 – 74 of *Systematic Zoology* and *Systematic Biology,* and used it to visualize changes in our field over time. The details of these and several additional analyses are provided in the Supplementary Information, which we hope will interest our readers. That said, those who only read this piece should still be able to comprehend our perspective on matters.

## Systematics

As we oriented our thoughts, we found we first needed to grapple with the scope of our discipline, bringing us face-to-face with the meaning of “systematics”. Our sense is that this has long been somewhat unclear or ill-defined, even in the minds of its avowed practitioners, and, based on our own experiences, it might have an especially nebulous meaning to outsiders and newcomers. Is systematics still “the scientific study of the kinds and diversity of organisms and of any and all relationships among them” (Simpson, 1961), or simply “the science of the diversity of organisms” (Mayr, 1969), or are these earlier definitions so broadly conceived that they end up including all of evolutionary biology, ecology, paleontology, etc. At the other extreme, most dictionary definitions – e.g., “the branch of biology that deals with classification and nomenclature” – feel entirely too narrow these days. Far better definitions have been proposed in recent years, one of which we’ll highlight soon enough. But, rather than adopting an existing definition, we tried instead to define systematic biology for ourselves, to solidify our views on the matter. So, who are we, more precisely?

It struck us that the meaning of systematics likely underwent at least two major shifts in the past 75 years: first, following the early rise of “phylogenetic biology” from the late 1960’s onwards, and again starting in the mid-1990’s as the field experienced a succession of advances, especially due to molecular approaches, and what these soon revealed about the nature of the “tree of life” itself. These shifts may have altered not only what we as systematists thought and did, but even what we published. So, we thought it might be useful to carry out an analysis of the literature. Using a pre-trained language model to embed scientific texts in a high-dimensional “article space” (Cohan et al. 2020; see Supplement for details), we found that similarity among the titles and abstracts of research articles published in *Systematic Zoology* and *Systematic Biology* tended to increase over time (Figure 1). Similarity among articles first rose around 1970, possibly triggered by debates among pheneticists, cladists, and more-traditional systematists. Similarity continued to rise for the next two decades, during a period when systematists published vigorously about cladistics and its consequences for biogeography and other fields. Similarity eventually plateaued entering the 1990s, possibly following the advent of “tree thinking” (O’Hara, 1988) and the new “comparative method” (Felsenstein, 1985), and not long before *Systematic Zoology* donned its new name in 1992. As we could not easily explain away these historical patterns as statistical artifacts (Supp. Figs. S1-2), we considered what they might indicate about our discipline? They could represent shifts only in writing style (not substance) across generations, or perhaps prolonged periods of jargon-riddled groupthink. As we interpret it, however, they represent distinct periods of alignment (and realignment) of systematics with phylogenetics.

**Figure 1.**
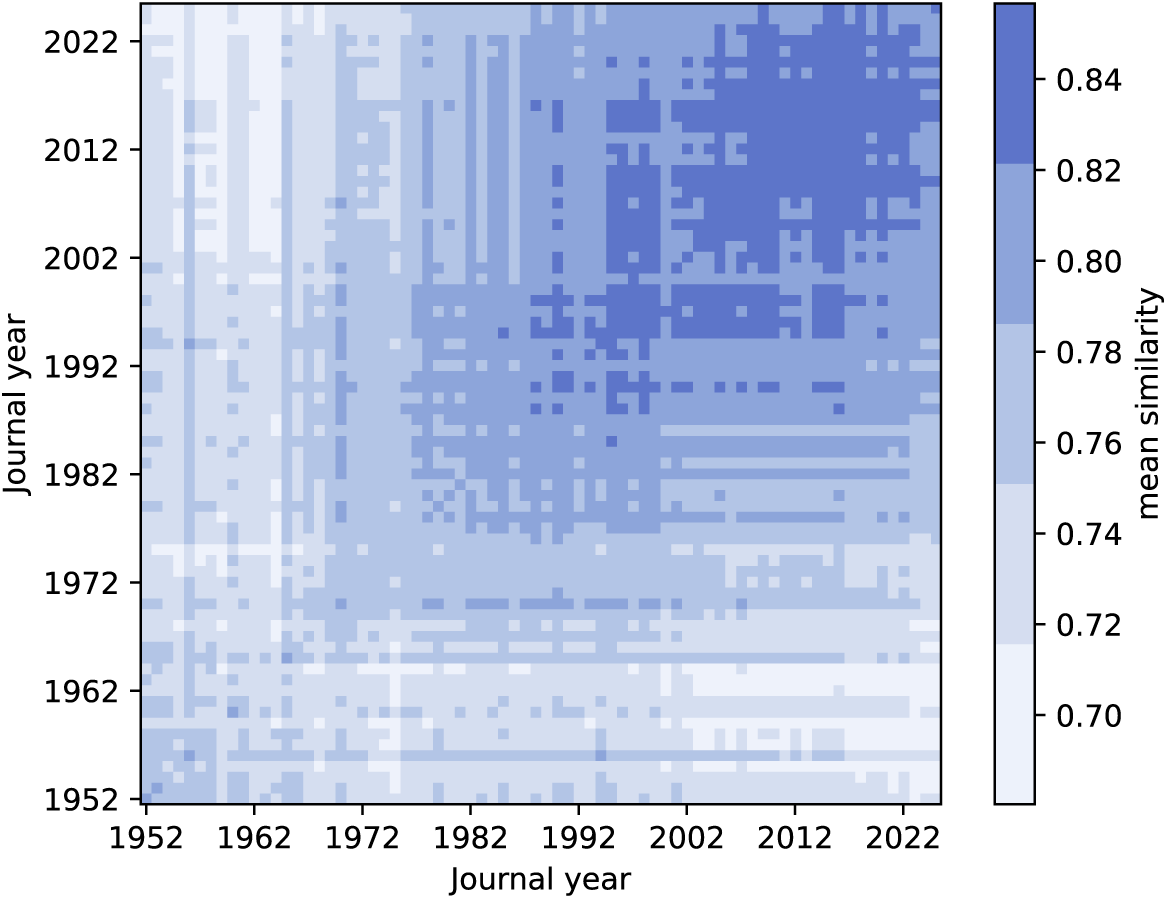
Mean similarity among research articles among different volumes of *Systematic Zoology/Biology.* Similarity scores may range from 0 to 1. See Supplement for details.

One possibility is that “systematic biology” has become nothing more nor less than “phylogenetic biology”. This would be convenient in the sense of making it easier to explain to our students and colleagues, and in this case maybe we should consider changing the name of our journal to “*Phylogenetic Biology*” and dropping the word systematics from our vocabulary altogether. But, are we then leaving out anything important? For example, as noted above, systematics has long been associated with classification and nomenclature. Are we still that discipline, or have we, in effect (and perhaps only inadvertently), shunted off some of these more traditional responsibilities? Maybe classificatory issues aren’t so much a part of modern phylogenetic biology, and instead they should be addressed in clade-specific journals? Spoiler alert: we think not!

More importantly, does “phylogenetic biology” actually capture the full range of interests of the people who have long associated themselves with systematics? One issue is how tightly phylogenetic biology is tied, conceptually and in practice, to the study of divaricating trees (the standard “tree of life” metaphor). Should we instead view it more broadly as encompassing evolutionary history, whatever form this takes – whether a tree, or a web, or something in between – along with all the genes, traits, and ecologies of the evolving entities? And, does “phylogenetic biology” really encompass the full range of motivations of systematic biologists, broadly speaking? We see our field as including not just people interested in developing phylogenetic methods and inferring phylogenetic relationships (these are, of course, prominently on display in *Systematic Biology* as we know it today), but also those who entered this arena enthralled by some particular group of organisms – who wanted to know everything there is to know about frogs, or mosses, or mushrooms, or whatever else. Donoghue and Edwards (2022) distinguished these two ends of the motivational spectrum as “phylogenetic biology” versus “clade biology”, and argued that both are necessary components of a healthy and more inclusive “systematic biology.” In effect, this may already have been recognized by the Editors of *Systematic Biology* with the distinction that was made (beginning in 2020) between regular “Research Articles” (“theoretical or empirical studies that explore principles and/or methods of systematics”) versus shorter “Spotlights” (that “focus on the focal group rather than the methodology”).

If *Systematic Biology* is the home journal for systematic biologists of all kinds, and if the health of the discipline depends on achieving a balance between these two halves, then how are we actually doing in terms of publishing? Using topic prediction (Grootendorst, 2022) to hierarchically group research articles from *Systematic Zoology/Biology* led to 39 topics that sorted into three main clusters (Figure 2; Supp. Figs. S3-5; see Supplement for details). Our interpretation is that the first cluster (Fig. 2, top) chiefly contains articles associated with classification, nomenclature, and phenetics, mostly representing the period before the 1990s. Papers in the second cluster (Fig. 2B, middle) predominantly concern empirical studies of particular groups of organisms through their traits and, later, through genomic data (cf. “clade biology”), mostly from the 1970s and onward. The third cluster (Fig. 2C, bottom) is home to most articles focusing on modern theory and methods (cf. “phylogenetic biology”), along with various topics that concern the technical aspects of phylogenomic analysis with some emphasis on applications to plants, from the 1990s through the present.

**Figure 2.**
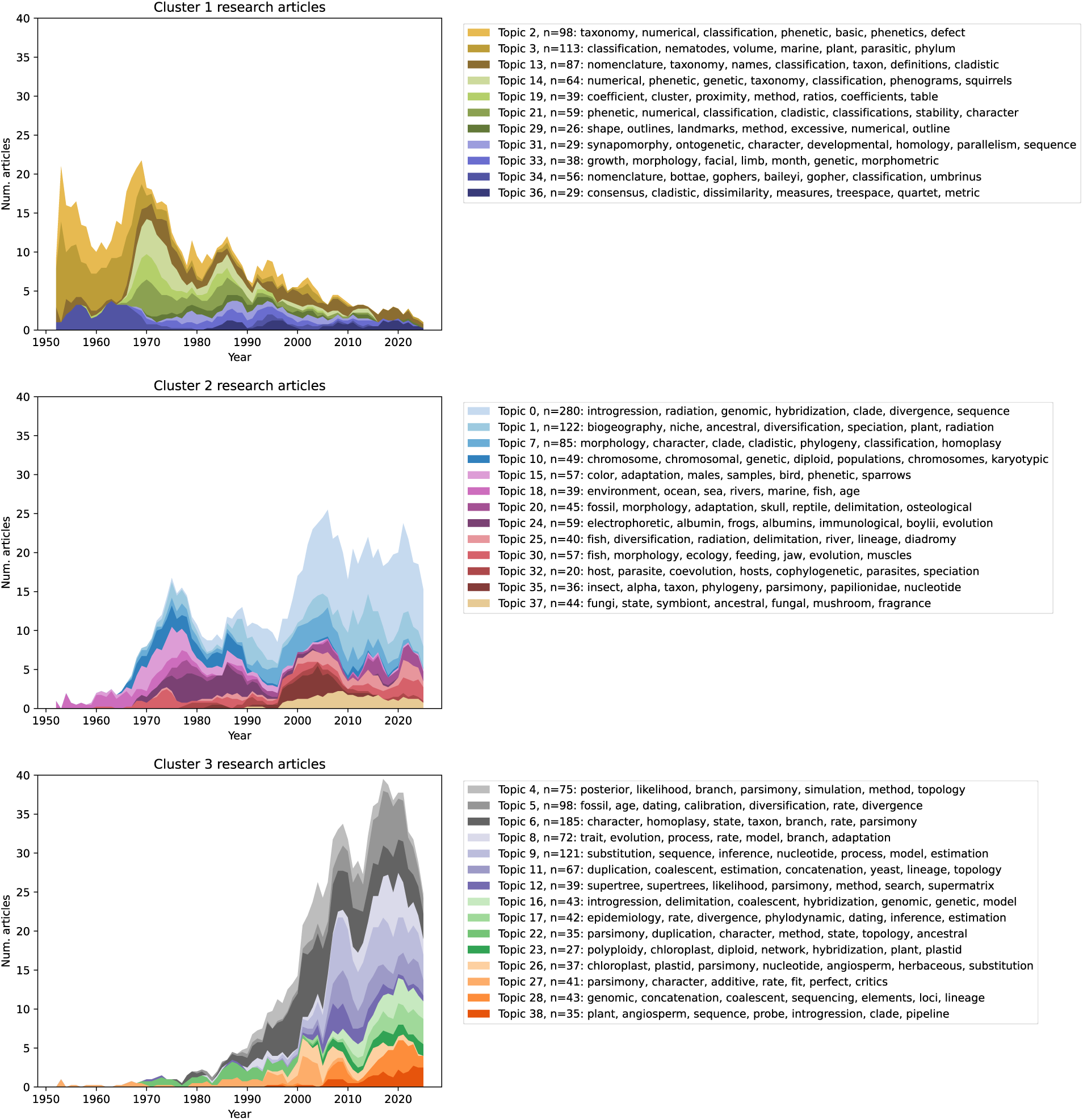
Numbers of research articles per topic published in *Systematic Zoology/Biology* during 1952 - 2025. Topic prediction divided articles into three main clusters, which then secondarily divided articles by inferred topic within each cluster (colors). Article counts were smoothed with a 5-year moving average to improve readability. Supplementary Figures S1-2 display the clustered topics. Supplementary Figure S3 shows the trajectories of all topics together, without being separated by cluster. See Supplement for more details.

We stress that the assignment of articles to topics, the labels associated with each topic, and the hierarchical clustering of topics all resulted from text analysis, but it is the two of us who have associated these clusters with the different elements of systematic biology. That said, we think that these three clusters are likely meaningful for three reasons. First, the predicted topics and their most-representative articles fell into clusters that are, to us, qualitatively appropriate. Second, very few articles (3.2%) were associated with multiple topics from different clusters. Third, topic prediction against a null dataset, with synthetic articles “bootstrapped” from the original pool of research article titles and abstracts, caused almost all (98.8%) articles to fall into one of just two topics.

Supposing that the clusters align with our interpretation, they would represent three, overlapping waves of topics, with Cluster 1 containing mostly older papers from *Systematic Zoology* (1952-1991), Cluster 3 (“phylogenetic biology”) containing mostly newer papers from *Systematic Biology* (1992-2025), and the enduring presence of Cluster 2 bridging the other two (1970-2025). Focusing only on articles published since 2000, citation rates for papers associated with “clade biology” (mean = 12.6 cite/yr, median = 8.7 cite/yr) and “phylogenetic biology” (mean = 16.4 cite/yr, median = 6.9 cite/yr) were similar (Supp. Figs. S6-7). That said, relative to “clade biology” papers, the top 1% of “phylogenetic biology” papers were cited more than twice as often and the bottom 1% were cited less than half as often, which could indicate that “clade biology” papers regularly find stable audiences, whereas “phylogenetic biology” papers more often experience “feast or famine” as the outcome. Lastly, while most topics in Cluster 2 and Cluster 3 align with “clade biology” and “phylogenetic biology”, respectively, topics in both clusters are blended with keywords from both ends of the spectrum, suggesting that modern “systematic biology” research is more integrated than it is polarized (Supp. Fig. S3).

That being said, what we have shown only applies to those “clade biology” and “phylogenetic biology” articles published in *Systematic Zoology* and *Systematic Biology.* Hull (1983; *pp.* 321-323) remarked that among the first 31 volumes of *Systematic Zoology*, the journal editors discouraged the publication of purely organismal or descriptive (“clade biology”) studies, and gave explicit preference to theoretical or conceptual (“phylogenetic biology”) studies. Based on personal experiences and conversations with colleagues, “clade biology” papers published in our journal tend to be more theoretical or conceptual than “phylogenetic biology” papers tend to be organismal or descriptive. Is it possible, for example, that the creation of Spotlight articles has had the unintended consequence of ghettoizing studies focused squarely on particular organisms, more than on methodology? Perhaps this is the balance that the readers, reviewers, and editors of the journal want, but perhaps not. Think about whether your favorite “clade biology” papers appeared in *Systematic Biology*, or were published elsewhere?

To us it seems clear that *Systematic Zoology* and *Systematic Biology* have provided a home for both “clade biologists” and “phylogenetic biologists”. One could argue about whether this is the right terminology, whether the predicted topics bear any meaning, or whether the current balance is just right or needs adjustment. In any case, we think it is important (at least from time to time) to take stock of the breadth of our discipline and examine how this is represented in our publications.

This returns us to the question we raised at the outset about the very meaning of “systematic biology”. Chris Simon reminded us (pers. comm.) that in 2001, when she became the Editor in Chief of *Systematic Biology*, she and several colleagues developed the definition of systematics that appears to this day on the SSB website (https://www.systbio.org/about-ssb.html):

**“Systematics** is the study of biological diversity and its origins. It focuses on understanding evolutionary relationships among organisms, species, higher taxa, or other biological entities, such as genes, and the evolution of the properties of taxa including intrinsic traits, ecological interactions, and geographic distributions. An important part of systematics is the development of methods for various aspects of phylogenetic inference and biological nomenclature/classification.”

This, we think, remains an excellent description of our field, and we hope that it has been (and will continue to be) widely consulted, especially by students who contemplate entering our field. At the same time, to us, the paragraph resembles the “2024 Word Cloud from the Members Meeting” displayed on the SSB home page (https://www.systbio.org; see Supp. Fig. S8), in the sense that anyone the least bit interested in this area of science can find a word or two that they identify with. But does this mean that they identify themselves with the field? Do they proudly declare themselves to be systematists, comfortable in the knowledge that others (including their parents) will know what they mean by this? We suspect not, at least judging by the students that we have queried, who typically prefer a more general label, such as “evolutionary biologist”, or a more specific one, such as “phylogenetic biologist” or “plant systematist“.

In our opinion, this definitional issue continues to need attention, and we wonder whether something snappy, perhaps along these lines, might be effective: *the study and representation of the shared evolutionary ancestry of biological diversity.* The two key elements here are fairly straightforward: “shared evolutionary ancestry” signifies both phylogenetic relationships and history, while “biological diversity” encompasses all forms of variation among lineages, taxa, species, traits, genes, and beyond. The word “representation” is included to reflect the classificatory objective – as Willi Hennig (1965) so aptly put it: “the expression of the results of . . . (our research) . . . in a form which cannot be misunderstood.” Whether or not this simplified definition helps, we urge continued consideration of the aims and scope of the science of systematic biology, so that we can clearly demarcate our domain and properly convey its fundamental importance to science and society.

## Collaboration

Over the past seven decades there has been a major transformation in how systematists operate. It used to be that students going into the field would choose a group of organisms to work on and become the sole expert on that group. One of the driving concerns was finding a group that nobody else was working on – there were plenty of clades to go around and, after all, having more than one person working on a clade would make taxonomic decisions that much harder. This is still a mode of operating but, for many of us, systematics has long since become a team sport. We think there are two main reasons for this. First, people have different interests and aptitudes, and it really helps to combine expertise – for example, a data-gathering organismal biologist might benefit from collaborating with a theoretician/methods developer, and vice versa. Given the pace of change and the time it takes to develop deep expertise of different sorts, it just makes good sense to join forces. Second, it is often just a lot more fun and rewarding to rub elbows with colleagues and grapple together with thorny issues. The two of us can attest to this in our own work together. And, we would add that we think there may be peculiar value in mixing scientific generations, bringing younger colleagues (often especially proficient in the latest methods) together with older ones (often especially knowledgeable about particular groups of organisms and the historical context of a problem). That said, it can be hard to find new collaborators from other traditions or eras of systematics. Maybe this calls for a matchmaking or “speed dating” analog for systematists?

Collaborations between countries are just as necessary and advantageous to systematics as collaborations between disciplines. For one, diverse perspectives and experiences produce an alloyed kind of research that is more resistant to scientific and cultural bias. In addition, the clades we study certainly do not pay attention to political borders, but we, unfortunately, need to. This can be a challenge, to be sure, but also presents us with opportunities to develop new models for international collaboration that defy the recent xenophobic tendencies of some of our home countries. Systematists simply cannot bow to isolationism, nor can we revert to the sort of scientific imperialism that characterized much of our activity over past centuries and decades. The exploitative practice of “helicopter systematics” must give way to the development of equitable and sustainable international field programs and partnerships (Ramírez-Castañeda et al., 2022). Forming these relationships can be difficult, but there are many useful avenues to pursue, from inclusive authorship, to student exchange programs, to the development of concrete benefit-sharing agreements with the communities in which we work (Kearney & Dittmar 2024). In any case, careful attention to the ethical foundations of our work must become the norm if we hope to continue our far-flung research efforts, and if we want to develop a globally activated biodiversity workforce. And, this we must do even in the face of misguided efforts in the United States and elsewhere to stymie efforts to promote diversity, equity, and inclusion. Our science, perhaps more than any other, absolutely depends upon it!

## Integration

Systematic biology has reinvented itself many times over, as new ideas and technologies have emerged, from evolutionary theory itself, to the discovery of DNA, plate tectonics, advanced imaging techniques, the computational revolution, and the genomic era. Take the rise of molecular phylogenetics starting in the 1980s, for example. Advances in genetic sequencing, mathematical modeling, and computational muscle ushered in unprecedented decades of discovery. Never before had we been able to infer the relationships of so many species in such exquisite detail – it has been a triumph beyond all reasonable expectations. Yet, if systematic biology is meant to explain how biodiversity emerged through shared evolutionary history, we need more – we need to thoroughly integrate population genetics, ecology, behavior, development, biogeography, and paleontology. But, also, depending on the question, we will need to connect with geologists, chemists, and physicists. In these mash-ups, systematists bring their rich knowledge of biological diversity and phylogeny to the table, and can often serve as ringleaders who cultivate fertile ground for testing ideas as these emerge across the sciences.

Systematics is, as botanist Lincoln Constance long ago emphasized, an “unending synthesis”, fueled by the continuous merging of diverse perspectives, new techniques, and intersecting disciplines (Constance, 1964). From our vantage point, there are three broad areas of integration that are especially likely to advance the field at this juncture.

First, integration of organismal biology into all aspects of modern systematics is key. It has become ever-easier to infer phylogenies, but doing so without that vital organismal context risks reducing systematics to “just” tree-building. Of course, tree-building is, in itself, a noble and onerous task. However, if, as we suppose, the primary goal of systematic biology is to deduce the histories of actual populations evolving and diversifying in and among ecosystems in space over time, then we also need deep knowledge of the structure and function of the organisms we are phylogenizing. Here we especially want to emphasize the word “deep”, to distinguish this type of organismal knowledge from simply consulting a database of well-known traits or the standard climatological layers, and mapping these onto a tree. Systematists (at least those at the clade biology end of the spectrum) should be digging deeper into their organisms, examining them closely and comprehensively, and making genuine new discoveries about them that can enrich our understanding of their evolutionary histories. Here, of course, there are obvious opportunities for collaboration with colleagues studying anatomy, physiology, development, functional genomics, etc.

Frequently, it is through the lens of organismal biology that we test whether our explanations for a particular phylogenetic pattern make sense or not. It is one thing to propose that, in general, polyploids have wider niches than their non-polyploid relatives, or that specialist parasites tend to cospeciate with their hosts, but it is a separate (and far more challenging) matter to measure those tendencies in clades of real organisms and to speak to their biological implications. Virtually all careful studies find gaps between the expectations of organismal experts and the results of data analysis. While there may be an impulse to hide these discrepancies as blemishes in otherwise perfect studies, making them apparent instead shines a light on the most fertile areas for future research: precisely where expert knowledge, theory, and data disagree. Keeping organismal biology centered in systematics also has cascading implications for who we train and how we train them, how we demonstrate the on-going necessity for our biological collections, and how broadly relevant we make our field through interdisciplinary collaborations.

Second, systematics must continue to integrate across scales. For example, within phylogenomics, certain characters and models are deemed more appropriate for resolving relationships at particular scales: from single nucleotide substitutions at shallower scales, to gene presence-absence and structural rearrangements at deeper scales. Where such divides sit usually depends on the underlying organismal biology of the clade under study (see above).

Analogously, in terms of taxonomic scale, we tend to analyze lineages at the population level to delimit species; at the species level for detailed analyses of smaller, densely sampled clades; at the level of backbone lineages to sort out overall relationships in larger, less densely sampled clades; or, at the broadest scale, incorporating all available data to build the largest, most densely sampled, and most comprehensive phylogenies possible. Analyses at these different scales often agree; the inferred relationships of backbone lineages may be the same whether one uses SNPs or amino acids or gene synteny, and may be largely unaffected by the exact sampling of tips. But we should pay special attention when more narrowly-focused and more broadly-focused analyses show discrepancies. Some of the most stimulating problems in systematic biology arise when these scales collide.

One thing that we think needs more attention is the relationship between detailed comprehensive analyses of small-ish clades, on the one hand, and massive but less comprehensively sampled phylogenetic analyses on the other. What is the peculiar value of studies of each type? And, how exactly do/can they inform one another? The development of “model clades” entails a deep dive into very many aspects of the organisms under study, and, consequently, these quite naturally engage colleagues from multiple disciplines (e.g., physiology, ecology, chemistry, etc.). As knowledge of phylogeny and biology grows ever stronger in these focal clades, they provide ready testbeds for hypotheses of all sorts, not to mention for methods development. Much broader studies are necessary to contextualize smaller clades but also especially in sampling multiple instances of various evolutionary phenomena of interest, thereby generating more statistical power for hypothesis testing. Our sense is that with a few notable model clade exceptions (e.g., oaks and sunflowers, Darwin’s finches*, Anolis* lizards, *Heliconius* butterflies), larger “mega-tree” analyses are often implicitly viewed as being more capable of producing more unexpected, revelatory, and/or generally applicable insights. We strongly doubt that this is true in general, and would argue instead that, as a field, we should be maintaining a healthy respect for, and balance of, studies of these different types.

Third, it’s crucial that we strive for integrative explanations for our findings – ones that not only allow, but welcome multiple causes. In storytelling, there is always a tension between simplicity and nuance: what details are necessary to explain a situation, and which ones are merely for color? This tension also looms when we describe evolutionary scenarios, explain tests for hypotheses, and even encode rules in our statistical models. And, we often resort (or are compelled by reviewers and editors) to communicating in highly telegraphed terms, which may then (at least inadvertently) mistake or misrepresent inferred simplicity for biological simplicity. For example, much of modern systematic biology relies upon statistical modeling to frame and test hypotheses. However, most models address only one or a few causal mechanisms rather than most of the valid ones, predisposing them towards simplified inferences from the outset. Also, complex multi-causal scenarios often require larger, high-quality datasets to generate statistical power, but such datasets are still rare. Simpler single-cause scenarios (think “key innovations” versus “synnovations”) that can be inferred from smaller datasets are therefore published more frequently. Lastly, and to no one’s surprise, simple scenarios are easier to communicate and digest. Consequently, we expect that the true complexity of evolutionary scenarios is underrepresented in conversation and in print.

Nonetheless, there are many remarkably clear and compelling studies finding evidence for complex, multi-causal scenarios. It is tempting to turn away from “tangled banks” in hopes of greener pastures – but, just as did Darwin, we must fully embrace and unpack the complexities of nature.

## Discourse

Any call for integration implies that there are rifts (or perhaps chasms) separating workers in their various subfields. Bringing differing viewpoints into contact naturally creates some friction, but, if managed well, this is highly likely to spark brand new ideas. Previous generations of debate in systematics featured passionate battles over the central ideas in the field: species concepts, homology, parsimony versus likelihood inference, and so on. This ferment surely fueled progress, but in some cases the friction touched off virtual “dumpster fires” that took years to die down. We are clearly engaged in a complex business, and disagreements are fully to be expected. The key to developing a vibrant discipline is to create the space for healthy, necessary debate, while avoiding the damaging vitriol that can emerge along the way.

With respect to debates, we think that one issue has simply been the growth of the field in terms of the numbers of researchers and the sheer volume of literature that we are creating. There are roughly 200 times more systematics papers published per year now as compared to the early 1980s (per Web of Knowledge), and thus the discourse is spread thinner than ever before. One of us (MJD) remembers fondly a time when it seemed possible to stay abreast of nearly all systematics papers as they appeared in print. Indeed, very conveniently, many of these were published in one place: in *Systematic Zoology* and *Systematic Biology*. With fewer papers being published, many researchers tended to actually read the majority of the papers in each issue of the journal, and they wrote directly and publicly to share their thoughts on the central issues that these raised. This helped to test the mettle of new ideas and celebrated collective engagement in what we do as systematists (Funk 2001). However, this direct, and often confrontational, form of communication also had its drawbacks. The other of us (MJL) heard harrowing tales of the “Dark Ages” (Felsenstein 2001), which ran from the earliest battles between the pheneticists and cladists, to the hostilities between proponents of parsimony versus likelihood-based approaches (Hull, 1988). This period enabled the formation of powerful cliques that controlled editorial decisions, enforced dogmatic ideas through the review process, and condoned or led to intellectual bullying. It might be argued that this accelerated progress, but these dynamics can, and actually do, have negative consequences, inhibiting the growth of valid (but contrary) ideas, favoring tribalism over scholarship, and endangering the careers of marginalized and/or junior scholars caught in the crossfire.

That said, comparing today to the past, our sense is that there is far less direct, public, and ongoing discussion in the literature concerning areas of friction in systematics, with some notable exceptions concerning how we model diversification or delimit species. Point of View articles in *Systematic Zoology/Biology* once served as the town square for publicly exploring ideas, but the number of POVs published per year has dropped off, while simultaneously increasing in length and technical complexity – in effect, they have become mini-research articles (Figure 3). Meanwhile, among research articles, it is common to read Methods and Discussion sections that appease both sides of a debate, rather than taking a stance on a controversial issue. Declaring a preference for one camp over others comes with some risk of backlash from reviewers and/or readers. In addition, empirical analyses are increasingly complicated, and require the use of dozens of hypothesis testing frameworks and methodologies. Formulating and defending a particular stance on every subject of debate would be exhausting but also impractical, for reasons of space alone.

**Figure 3.**
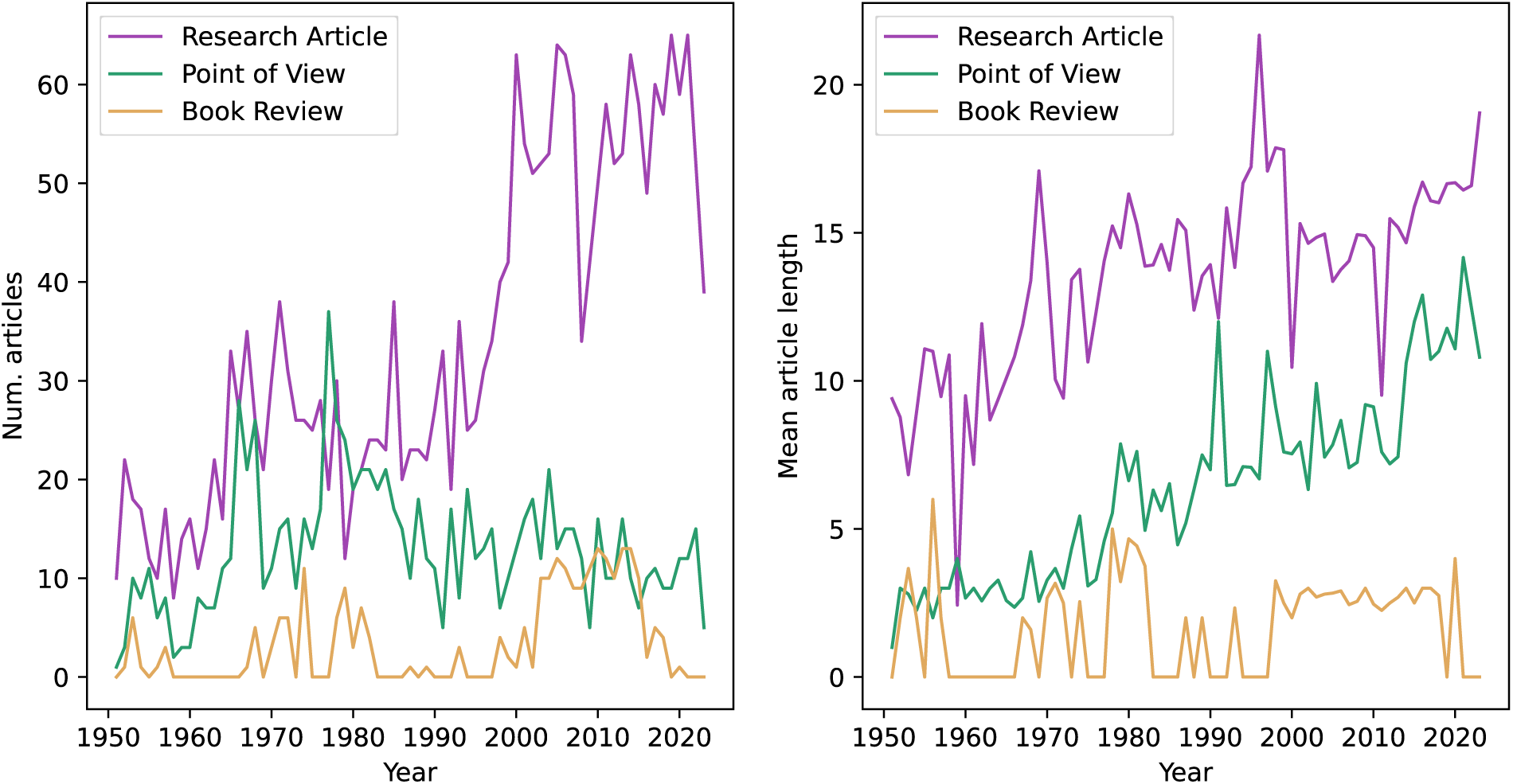
Number of articles (left) and mean article lengths (right) published in *Systematic Zoology/Biology* during 1952 - 2025. Articles are separated into Research Articles (purple), Points of View (green), and Book Reviews (gold). Other article types are not shown. Article lengths are measured in numbers of pages, rounding up. See Supplement for more details.

What should the Society of Systematic Biologists (SSB) and our publications be doing to ensure that systematics has an active, engaged, and yet professional culture of discourse going into the future? The debates and panel sessions held in recent years at the SSB standalone meetings have been stimulating, and have helped to nurture a healthy culture of discourse.

These sessions elevate issues that matter to systematists (e.g., inferring extinction from extant taxa, phylogenetic resolution, diversifying participation in systematics, etc.), create space for them to be discussed equitably, and to model good behavior towards our colleagues, even those we might strongly disagree with. One downside to these sessions is that they are in-person only, and even still may be a bit too large to promote a level of engagement that could significantly advance understanding of contentious topics. With only a little effort, we can imagine that *Systematic Biology* could help choreograph finer-scale engagement of this kind within the community. For example, specifically inviting short commentaries on recently published work, as other journals do, could benefit *Systematic Biology* and strengthen a sense of community in its readership. Another option would be to invite authors to write separate, opposing views, or attempt to write a single, synthetic view to highlight a currently unsettled issue (e.g., Platnick and Harper, 1978; Felsenstein and Sober, 1986). A third idea would be to host virtual “parlor sessions” following each issue of *Systematic Biology*, where authors and readers could discuss what they found most interesting or concerning. Lastly, we see immense value in reincarnating true Point of View articles that succinctly and respectfully raise open-ended questions for consideration by the community.

## Infrastructure

Traditionally, the identity and the day-to-day practice of systematics has been intimately connected with natural history collections, and this makes good sense. Collecting and curating are deeply part of the mindset of many who have entered systematics over these decades (though perhaps less so toward the present?). Not only that, the documentation and naming of biodiversity and the data that we depend upon to do our work ultimately traces to museum collections and their associated libraries. Given that this is true, it behooves us as a discipline and as a scientific society to ensure that this crucial infrastructure is thriving – that museum collections are actively developed, curated, and safely maintained to support research, teaching, and outreach by all. Many of us have taken the existence of such collections for granted, but this is clearly a mistake. Recent high-profile divestments (e.g., the undoing of the Duke herbarium) and the withdrawal of federal funding for programs that support museums (e.g., the Institute of Museum and Library Services, IMLS) should be a wake-up call, and we, as a scientific society, need to intervene constructively, and in some cases very quickly, to protect these resources. The threat is clear, but it is less obvious how we can be most effective in safeguarding these key resources. Adjacent organism-oriented societies (e.g., the American Society of Plant Taxonomists, the American Society of Ichthyologists and Herpetologists) will surely play a role, but, in our opinion, this is not a responsibility that we should leave totally to others. This issue warrants our immediate attention.

Beyond physical specimens, our community is responsible for a wide range of digital resources that power our research and the real-world applications of biodiversity knowledge (e.g., through the Intergovernmental Science-Policy Platform on Biodiversity and Ecosystem Services, IPBES). We have spearheaded global efforts to digitize museum specimens and extract label data on geographic locations, phenology, etc. Facilities of various sorts have emerged to generate and provide access to these data (e.g., The Global Biodiversity Information Facility, GBIF; Integrated Digitized Collections, iDigBio), but these too will need to be supported and protected against short-sighted funding cuts. The same thinking applies to the repositories of genetic, genomic, and transcriptomic data, and the scientific software packages that our community both produces and uses so prodigiously. It is these kinds of resources, collectively, that allow society to identify and respond to threats such as habitat destruction, invasive species, and disease outbreaks. For example, it was through decades of sustained investment in this infrastructure that allowed scientists to quickly determine that COVID-19 was not merely “a bug”, but a newly emergent coronavirus, bringing vital phylogenetic context to how we named and tracked various strains throughout the pandemic. With major science cuts currently on the docket in our home country’s funding agencies, including to the National Institutes of Health (NIH) and the National Science Foundation (NSF), we are reminded that we must all work actively to ensure that such resources persist and provide open access to the data that we depend upon, regardless of governmental support. And, there are other repositories that need our attention. For example, what can we do as a community that generates phylogenetic knowledge to provide better long-term storage and access to phylogenetic trees (think TreeBASE, Dryad, etc.), underlying molecular and morphological datasets, and scientific software, which are very often the primary and longest-lasting products of our research efforts?

We also consider classification systems and taxonomic names to be critical infrastructure that our systematics community provides to science and to the world at large (e.g., through Wikipedia). These are crucial in orienting research and information retrieval and, in this respect, are even more influential in the age of AI. In our opinion, SSB should be deeply involved in training its membership to use newer machine learning techniques critically and responsibly, otherwise centuries of domain expertise cultivated by our field could risk being overshadowed by sloppy AI-assisted work or replaced by unvetted AI-based automation. Instead, systematists properly trained to use AI could revolutionize how we generate, represent, and disseminate high-quality phylogenetic knowledge.

It is therefore vital that *Systematic Biology* continues to publish work that advances our thinking on general theoretical issues associated with classification and nomenclature (e.g., see discussions of “phylogenetic nomenclature” and the *PhyloCode*). Moreover, we should encourage authors, in whatever ways we can, to go-the-distance and actually translate their phylogenetic findings into new classification systems, creating new names as necessary to unmistakably reflect evolutionary history. Sadly, we so often stop short of providing this truly enormous added value. To incentivize this important work, one possibility would be to develop a special index to properly credit systematists for naming clades; for example, associated with entries in the RegNum database that manages the registration of names under the *PhyloCode*. Perhaps SSB could even sponsor “name-athons” (cf. “hackathons”) at our annual meetings to provide expert guidance on the sometimes daunting details associated with formal nomenclature under any/all of the various nomenclatural codes.

## Society

Here we are, writing an article for a society-run journal, and here you are, reading it. Would this journal be what it is today had it not been backed by a scientific society? After all, no one really needs a “home journal” – why not just publish anywhere and find articles as needed? This attitude might serve individual needs, in the moment, but not the interests of a field over generations. Throughout most of the last century, journals were primarily run by societies in service of advancing their discipline. To this end, each society helped meet its own journals’ needs, including the recruitment of editors, management of finances, working with publishers to print and disseminate physical copies of the journal, and setting the standard of rigor for its articles. In contrast, for-profit journals, run without society involvement, put financial interests above the community. It’s no secret that for-profit journals routinely pressure authors and editors to prioritize quantity over quality, but in service of whom?

Another possibility would be to boycott subscription journals all together, society-run or otherwise, and skip the peer-review system, too. Instead, we could work through non-profit publishers such as *arXiv* or *bioRxiv*, where one undoubtedly can find diamonds in the rough, even though there’s a lot of rough. Alternatively, *Systematic Biology* and *Bulletin of the Society of Systematic Biologists* are run by systematists for the systematics community. Articles in these journals are consistently of high quality and relevance, documenting what we as systematists think and do as a community. Of the various options, we strongly favor this one.

If it is fair to ask what purpose society-run journals serve, it is also fair to ask what purpose scientific societies themselves serve beyond their journals. Ultimately the question becomes: why should one join SSB, or any scientific society, and pay the dues? Fundamentally, scientific societies bring people together around a shared scientific interest. In the case of SSB, we assume that we are all, on one level or another, fascinated by the improbable, intricate, and endlessly weird evolutionary history shared by all living things. Many of us are also deeply concerned with the fate of biodiversity at the hands of humans. It seems important to have a platform to band together around these interests. But, there’s more to it than that. Historically, SSB has a strong track record for supporting its community. One measure of this is through awards, from Graduate Student Research Awards for junior scholars, to the IDEA Award (with the American Society of Naturalists and the Society for the Study of Evolution) for promoting diversity and inclusivity in the sciences, to the President’s Award in recognition of systematists who have dedicated their lives to our discipline. Another measure is the tireless work of co-organizing the annual joint Evolution conference (again, with ASN and SSE) and our biannual standalone meetings. And then there’s the development of new diversity and mentorship programs, and advocating on behalf of the interests of systematists to the government, other relevant agencies, and to the public. Obviously, this last function has now taken on a special urgency, with many scientific institutions and funding agencies around the world actively under attack and evidently broadly misunderstood by politicians and the public at large. Whether you are simply a dues-paying member, a volunteer council member, or an editor, everything SSB does is made possible by the generosity of you and your colleagues. We are organized bottom-up by community members, not top-down by institutions. Get involved, stay involved, and make SSB even better!

Just as the practice of systematics changes over time, so must our society adapt to changing circumstances. Reimagining SSZ as SSB opened the door to thousands more systematists, solely by expanding the breadth of our taxonomic focus. Recent suggestions that we reimagine our society again as the Society for Systematics and Biodiversity might better reflect how our core community views the society, while casting an even wider net. SSB has also expanded itself geographically by (co-)hosting conferences in countries including Brazil, Canada, France, and Mexico, offering Evolution conference registration discounts through the Global Membership Assistance Program, and electing more and more international colleagues into leadership positions. These efforts are commendable, but are only beginning. We sincerely hope that SSB will become a truly global society and an even greater force for good.

## Conclusions

Returning to our opening thoughts, what phase is systematic biology in now? In our view, we very definitely experienced a “scientific revolution” with the dramatic shift into the phylogenetics era from the 1960’s to the 1990’s. In the relatively “normal” phase that has followed, our empirical studies have fleshed out – clade by clade, and sometimes in stunning detail – the Tree of Life. But as these studies proceeded, they rather soon revealed complexities that then touched off another major conceptual shift, this time re-envisioning our fundamental notion of “the tree”.

By the early 2000s, systematists flocked to statistical phylogenetics for practical and philosophical reasons, replacing the quest to infer “the tree” with inferring distributions of highly probable trees. By embracing historical uncertainty as a fact of life, rather than a corroding substance, systematists innovated increasingly realistic models that were better suited to testing complex evolutionary hypotheses. In parallel, the “gene trees within species trees” realization (see especially Maddison, 1997) set off scores of new methodological developments and empirical discoveries centered around gene coalescence, incomplete lineage sorting, and admixture on various scales. These are issues that we are clearly still grappling with today, with ramifications for multiple areas of inquiry, from adaptation and biogeography to coevolution and community ecology. There have, of course, been other major developments that bear on systematics and its adjacent areas, most importantly the availability and integration of detailed geographic locality and morphological data from museum collections to better infer geographic ranges, climatic niches, and phenotypic evolution, and how these have changed through time.

It is hard to predict where all of this will lead, but we find ourselves in a very exciting period in which many of the once-distinct boundaries between micro- and macroevolution – and even between ecology and evolution – have blurred if not vanished. Over the past twenty-five years, “tree-thinking” has played an increasingly prominent role in ecology, genomics, developmental biology, medicine, and microbiology, and perhaps these are next to unify with systematic biology. In this new space we anticipate a return, with fresh eyes, to the age-old problems of our discipline: determining the nature of species, homology, diversification and such. Will we “solve” these problems this time around? Maybe not, but we will have fun trying, and, as always, we will surely deepen our understanding of the living world in the process.

One last thought. It is vital for our community to periodically reflect, as we are now, upon what systematic biology has experienced through its lifetime, and what has caused it to take the shape that it has. The 75th volume of *Systematic Biology* is slated to contain special articles that consider the history of our journal and of our discipline more generally. Read these and learn from the past. Browse through earlier volumes, and pause to consider what questions our forebearers pondered. Not just the questions they faced as systematists, but also the questions they faced concerning the relevance of systematics and science to the broader world. We encourage budding systematists to get in touch with older colleagues to grapple with fundamental issues in the field. Likewise, long-established systematists should introduce themselves to younger colleagues and find out what’s on their minds. As we ourselves have discovered, one excellent starting point might be: what exactly is systematic biology these days? Great ideas and collaborations are bound to follow!

## Acknowledgements

We thank Chris Simon, in her role as the Chair of the SSB Legacy Committee, for inviting us to contribute the first in a series of special papers for the 75^th^ volume of SSB. Many thanks also go to Bob Thomson, editor-in-chief of the journal, and Andy Seagram, with the journal’s publisher at Oxford University Press, for providing data tables of past articles published in *Systematic Biology* and *Systematic Zoology*. Finally, we are grateful to Erika Edwards, Chris Simon, Bob Thomson, and two anonymous reviewers for their very helpful comments on earlier versions of our manuscript.

## Supplementary Information

Our supplementary document describes how we assembled our dataset and explains how we analyzed journal articles. It also contains additional analyses and figures relevant to the interpretation of our results.

## Data Availability Statement

Our dataset and analysis code are hosted at https://github.com/mlandis/systbiol75. We have also archived a copy of this repository on Data Dryad: http://datadryad.org/share/E10TCz9N8_5exW-IJsE63Un2806TZtx_ZXh2szdb7Go (reviewer link).

