## Supplementary Information for "Seventy-five Years of Systematic Biology: Looking Back, Moving Forward"

### Overview

Datasets, code, and output we produced for this paper are hosted on GitHub: https://github.com/mlandis/systbiol75. Files are also archived on DataDryad: https://doi.org/10.5061/dryad.905qfttz8. Code is written in Python and shared as an interactive Jupyter notebook.

### Dataset

We built a table containing 5,150 published records from *Systematic Zoology* (*SZ*) and *Systematic Biology* (*SB*). This table was last updated in June 2025, so it does not include all records that will appear in Volume 74 (2025), and it includes no records for Volume 75 (2026). Volumes 1 – 40 for *SZ* correspond to years 1952-1991, and Volumes 41 – 74 for *SB* correspond to years 1992-2025. Volumes 1 – 49 each generally contain 4 Issues, whereas Volumes 50 – 74 each have 6 Issues.

We built our dataset in several stages, beginning with a table provided by Andy Seagram at Oxford University Press, kindly arranged by Bob Thomson and Chris Simon. This table was generated in May 2025 from Dimensions AI (https://www.dimensions.ai/) database records and contained information for 4,655 records associated with *SZ* and *SB* between 1952 and 2025. The Dimensions table included five important fields of interest to us (article title, publication year, volume, issue, and DOI) that were almost perfectly complete and consistently formatted, making them suitable for statistical analysis. Three other fields of interest (article abstract, article type, and citation count) were variously incomplete, inconsistently formatted, and/or inaccurate, which each required different correction strategies to amend.

First, the article abstracts presented two kinds of problems. Among research articles, 293 of 2,757 records in the Dimensions table lacked entries for the abstract field. Initially, we thought this was a defect with the Dimensions table, but it appears that almost no articles before 1967 contained Abstract sections. Among research articles published in 1967 and onwards, 203 research articles lacked text in the abstract field. Because the content of research articles from early volumes of *SZ* and *SB* were only available online (https://academic.oup.com/sysbio/issue) as documents (.pdf) that must be individually downloaded and manually processed, we could not extract abstract information for large numbers of documents automatically. We decided not to update the abstracts for these articles. A second issue is that most abstract fields for articles associated with (approximately) Volumes 18 – 31 also include the article title, authors, and publication information, which could potentially cause dissimilar papers to appear spuriously similar due to shared metadata. For this reason, we removed these extraneous metadata from all abstracts.

Second, the original table contained different citation counts from Clarivate (through Web of Science) and Dimensions AI, but many of these records lacked citation counts or vastly undercounted the numbers of citations reported by Google Scholar (e.g., 10 versus 1,000 citations in some cases). Because Google Scholar does not allow bulk retrieval of article citation, we used PublishOrPerish (Harzing, 2007) to generate citation counts for all articles associated with *SZ* and *SB* in 10-year batches. We used ISSN 0039-7989 for *SZ* and ISSN 1063-5157 for *SB* to minimize the chance that our search collected articles from other sources that had a partial match with the journal name. In cases when Google Scholar associated multiple records with a single article, we only retained the record with the highest number of citations. We used article titles to cross-reference the datasets and add Google Scholar citation counts to our main table, and manually updated the citations for any records that had no obvious match. Google Scholar data were collected in June 2025.

Lastly, article type fields were often imprecise or inaccurate. The Dimensions dataset only classified articles using generic groups (e.g., articles, book reviews, editorials) and not the custom types used by *SZ* and *SB* (e.g., research articles, points of view, spotlight articles, symposium articles, etc.). However, articles hosted on the *SB* website were correctly sorted into sections that matched the custom types we desired. We wrote a Python script to scrape each article's type along with its title, abstract, DOI, page count, and other relevant information from all issues of *SZ* and *SB* that were hosted on the *SB* website. This netted 5,095 scraped articles in total. We then updated the article types in our main table by matching DOIs and/or titles between the tables. Our new dataset scraped from the *SB* website contained 496 articles not contained in our main table, which we added and populated manually.

As part of building this dataset, we also skimmed through the entire dataset several times to check for general correctness. In cases when we noticed individual records or fields that seemed incorrect, we updated them either by visiting the *SB* website, Google Scholar, or downloading the article in question. After including all verified journal articles, removing duplicate entries, and resolving inaccurate article types, our new table contained 5,150 articles with 2,531 research articles, 964 points of view, 326 book reviews, and 1,329 other miscellaneous articles (e.g., announcements, spotlight articles, symposia articles, etc.). Although we believe this table represents the most accurate record of *SZ* and *SB* articles in current existence, we have no doubt that it still contains undetected errors and missing entries/fields.

### Analyses

*Article similarity through time.* We measured the similarity among 2,531 research articles published across different volumes of *SZ* and *SB*. To do so, we concatenated the titles and (when present) abstracts of our articles, then passed those strings (the texts) into a pre-trained sentence transformer, SPECTER (Cohan et al., 2020) to embed each article into a 784-dimensional vector space. SPECTER was trained using the titles and abstracts of roughly 146,000 scientific articles from Semantic Scholar (Ammar et al., 2018), making it well-suited our purposes. Using the embedded coordinates for the articles, we then computed the mean cosine similarity between all pairs of articles in each pair of volumes. We chose cosine similarity because it only measures the degree to which two embedded vectors are aligned, but ignores the magnitude of the similarity, which could be influenced by the fact that some texts only contained a title (e.g., before 1964) whereas others contained a title and abstract. A cosine similarity score of 0 means the embeddings are orthogonal in vector space and a score of 1 means the directions of the embedded vectors are identical. We then plotted the similarity scores as a heatmap (Figure 1 in main text).

To assess whether any changes in similarity among SZ and *SB* articles through time was of statistical interest, we compared the scores in our main dataset (*D1*) described above to scores for three alternative datasets (*D2* – *D4*). We first constructed an alternative dataset (*D2*) from the same *SZ* and *SB* research articles as in *D1*, except we shuffled the publication years among volumes, allowing us to test whether it also induced apparent shifts in similarity. We then generated two more datasets by randomly sampling 20 research articles from the SciDocs dataset used to train SPECTER for each publication year during 1970 – 2019 (SciDocs contained too few records outside of these dates). The first of these two datasets (*D3*) retained the original publication years, whereas the second dataset (*D4*) shuffled publication years among articles (as above). These datasets allowed us to assess the background levels of similarity among all research articles, regardless of discipline, with and without temporal information. For all four datasets (*D1* – *D4*), we generated similarity score heatmaps analogous to Figure 1 and plotted the within-year similarity score across all four datasets to aid comparison. Only *D1* demonstrates a clear pattern of increased similarity among articles over time, both for the heatmap of between-year similarities (Supp. Fig. S1) and the line plot of within-year similarities (Supp. Fig. S2). We interpret these results as support for increased similarity among *SZ*/*SB* research articles published near 1970 and 1990; these increases are not easily explained as random noise or caused by academia-wide changes in writing style.

*Topic modeling.* We used BERTopic (Grootendorst, 2022) to predict interpretable topics for our embedded texts. BERTopic takes a set of embedded texts, clusters the embedded texts by similarity, and then associates all clusters with representative keywords to construct human-interpretable topics. The number of topics, the exact topic keywords, and assignments of texts to topics are treated as unknowns to be learned from the texts by the analysis.

BERTopic has six main steps: embedding, dimension reduction, clustering, vectorizing, keyword discovery, and representation. The first three steps (embedding, dimension reduction, and clustering) determine which articles are assigned to which clusters. The last three steps (vectorization, keyword discovery, representation) determine exactly which keywords are used to represent each cluster. We used the same text embeddings that were generated for Figure 1. For dimension reduction, we used UMAP with Euclidean distances, a neighborhood size of 20, and a minimum distance to reduce dimensionality to represent our embeddings with 5 components. Clustering used HBDScan with Euclidean distances, minimum cluster sizes of 15 and minimum sample sizes of 3, using the excess of mass (eom) criterion for cluster selection. For vectorization, we used a standard count vectorizer to convert the texts into tables of n-gram frequencies (1 to 3 words) while removing common English stop words. Keyword discovery then used categorical Term-Frequency Inverse-Document-Frequency (c-TF-IDF) to find sets of keywords that were relatively unique to each topic, while providing a list of 132 terms from systematic biology as suggested seed words for topics with an up-weighting factor of 2, and using the Okapi BM25 ranking function to boost the importance of rare terms. To improve the final representation of keywords for each topic, we used both the Maximum Marginal Relevance model with a diversity score of 0.3 (mild diversity enhancement) and the Parts of Speech (POS) model from spaCy with the *en_core_web_lg* trained pipeline to eliminate different grammatical forms of the same word (e.g., fish vs. fishes). Lastly, we used built-in tools from BERTopic to construct a topic hierarchy, using 1 minus the cosine similarity for the distance metric. These settings were chosen based on best practices recommended by the BERTopic documentation, familiarity with the dataset, and trial-and-error.

Our analysis resulted in 1,799 texts being assigned to 39 topics with high probability. We inspected the summary of topics and representative texts to validate the topic predictions were sensible. Next, we considered the 732 texts that were left uncategorized. We assigned each of these texts to the topic with the highest probability, while leaving alone texts with >0.99 probability of being uncategorized. This resulted in all 2,531 texts being assigned predicted topics. Figure S3 displays the topic assignments of all research articles in a two-dimensional space. The "topics_info.csv" and "texts_info.csv" files provide information on topics, topic keywords, research articles (texts), and topic assignments. We used these files to assess the topic assignment quality.

We visualized the hierarchical topic scheme using modified versions of BERTopic plotting tools (Figure S4). We then extracted the subtopics associated with the three deepest clusters ("clades") of topics. For this analysis, the first cluster (top) generally contained topics with keywords associated with classification, nomenclature, and phenetics. The second cluster (middle) primarily contained topics associated with specific clades, organismal traits, or genomic analysis. The third cluster (bottom) contained topics with keywords concerning phylogenetic methods and/or concepts, along with a group of topics regarding the use of phylogenomic models in plants. The representative papers of topics associated with phylogenomics in plants in Cluster 3 have titles such as “Mitochondrial Phylogenomics Of Early Land Plants: Mitigating the Effects of Saturation, Compositional Heterogeneity, and Codon-Usage Bias” (Topic 27) and “A Universal Probe Set for Targeted Sequencing of 353 Nuclear Genes from any Flowering Plant Designed Using K-medoids Clustering”, and have abstracts that place far more stress on methodological innovations over the organismal research. For these reasons, we associate the first cluster with traditional concepts in systematics, the second cluster with “clade biology”, and the third cluster with “phylogenetic biology” (see main text for discussion of “clade biology” and “phylogenetic biology”).

We stress that while the topic names were generated by the pipeline and can be considered as results from data analysis, the cluster names were assigned by us (the authors) and should be considered interpretations. It may be that the predicted topics, the clusters, and our interpretation of the clusters (i.e., as representing “clade biology” and “phylogenetic biology”) are all meaningless. To consider this possibility, we inspected all predicted topics, their keywords, the titles of their most-representative articles, and several random articles associated with each topic to gauge whether the prediction was reasonable. We additionally reviewed which topics were associated with which clusters. We judged that the topics within each cluster seemed more similar with each other than with topics from other clusters, and that the topics and most-representative articles associated with each cluster were reasonably accurate (i.e., there was far more signal than noise).

We performed two analytical tests to determine whether the topic prediction pipeline was prone to producing meaningless topics and clustering schemes. First, we measured the proportion of "multi-topic" articles that were associated with probability >0.05 in each of two or more topics, and where the second-best topic received at least half as much support as the first-best topic. We identified 965 (38.1%) "multi-topic" articles in our dataset, of which 885 (35.0%) were associated with only one cluster and the remaining 80 (3.2%) were associated with different clusters. This is expected, since the clusters correspond to the deeper, hierarchical signal in similarity among topics. Second, we generated a dataset of topic-free research articles, where each article was represented by one random abstract and 5-6 random sentences sampled (with replacement) from the original *SZ/SB* research articles; that is, no article was expected to have a cohesive topic. We then re-ran the topic prediction pipeline and found that 2,502 of 2,531 (98.8%) of the randomly constructed articles were placed into one of two topics, with both topics falling into a single cluster. This suggests that randomly generated *SZ/SB* research articles fail to produce meaningful numbers of topics or clustering hierarchies, unlike what we obtain from the actual dataset.

To visualize topics through time (Figure 2 in main text), we collected the counts of articles associated with each of the three main clusters and each topic for each year. Because the number of articles within a topic could fluctuate radically from year to year, we used a 5-year rolling average for article counts to make the plot easier to read and interpret. Whereas Figure 2 separates the visualization of topic dynamics through time by cluster membership, Supplemental Figure S5 displays all topics together, ignoring the clustering scheme.

We note that BERTopic is a stochastic method with a wide variety of tuning parameters. The exact composition of predicted topics, their keywords, and their hierarchy can be somewhat sensitive to analysis settings. However, in general, our experimentation with topic modeling typically led to three main clusters, each composed by similar topics as presented in Figure S4.

*Article types through time.* We summarized article types through time in two ways (Figure 3 in main text), as the number of articles per type and the mean lengths of articles per type for each year. We did not perform any statistical tests for this visualization.

*Citations by type through time.* We explored whether articles associated with the “clade biology” or “phylogenetic biology” clusters had different citation patterns. Articles concerning “clade biology” tend to be of greatest interest to other experts working on the group, and potentially suffer from lower citation rates due to being overspecialized, taxonomically speaking. On the other hand, an excellent “clade biology” study can become the emblem of an evolutionary process and/or provide an ideal dataset for secondary analyses. That said, many “phylogenetic biology” articles describe new models, techniques, or software that become immensely popular in the field, while others can be exceedingly technical or abstract and have no apparent relevance to practicing biologists. Of all these options, we assumed that extremely popular software packages associated with “phylogenetic biology” would lead to higher mean citation rates when compared to “clade biology”. To account for the fact that older articles generally have had longer periods of opportunity to accumulate citations, we compared articles on the basis of citations per year, computed as the number of total citations divided by the publication age (time since publication + 1). In addition, because articles published within the past 20 years tend to have a higher citation rate than older articles, we limited our comparison to the period of 2000 through 2024 (Supp. Fig. S6).

Recent research articles associated with the "phylogenetic biology" cluster indeed tend to be more highly cited (mean=16.4) than "clade biology" articles (mean=12.6), with papers associated with "traditional systematics" being cited the least often (mean=8.2; Supp. Fig. S7). However, "clade biology" papers have lower variability in citation rates (sd=18.4) than "phylogenetic biology" articles (sd=64.3). When using median citation rates, which are less sensitive to extreme values, "clade biology" papers (median=8.7) instead appear to be cited more often than "phylogenetic biology" papers (median=6.9). The most highly cited "phylogenetic biology" papers ($99^{th}$ quantile=139.9) tend to be cited more than twice as often as those from "clade biology" ($99^{th}$ quantile=59.1). Our interpretation is that while a few "phylogenetic biology" articles, particularly software methods, may "strike it rich" and become widely cited, "clade biology" and "phylogenetic biology" papers are cited at similar rates, on average.

### Reflections

David Hull performed a variety of bibliometric analyses in *Science as a Process* (Hull 1988) by individually and manually processing the articles available at that time. This included classifying papers into various categories (e.g., "Pro-Phenetics versus Anti-Phenetics") based on article content. Newer automated language processing techniques, such as those we used in this study, could broaden and complement the original work of Hull (1988).

A practical challenge is that automated language analysis requires that the data are complete, consistently formatted, and accurate. Producing a dataset with these qualities was more challenging than we had originally anticipated for several reasons. First, information about the journal itself is scattered across several proprietary and/or private databases. Second, many records in those databases are inaccurate and/or in conflict, making it difficult to know what records can be trusted. Lastly, because the journal itself has changed substantially over time (e.g., in terms of journal name, article types, article formatting, and management), it is difficult to draw comparisons across volumes even if the content is accurate.

The Society of Systematic Biologists (SSB) should consider maintaining its own table for all articles published in *SZ* and *SB*. Oxford appears to have some ability to improve the quality of records it provides us. However, ultimately, we think that curating data for the ideal table will require a combination of automated and manual data processing. SSB Legacy Committee Chair, Prof. Chris Simon, raised the idea that we (the authors) gather a team to help with this work. While we did not have enough time to recruit and train others, it seems like an excellent idea for the future to ensure that SSB has an accurate and handy record of what articles have appeared in its flagship journal as well as for the *Bulletin of the Society of Systematic Biologists*.

For anyone who has read this far, we encourage you to download the table of *Systematic Zoology* and *Systematic Biology* articles and take the time to witness and appreciate the extraordinary diversity of research our community has produced these past 75 years!

### *Supplementary Figures*

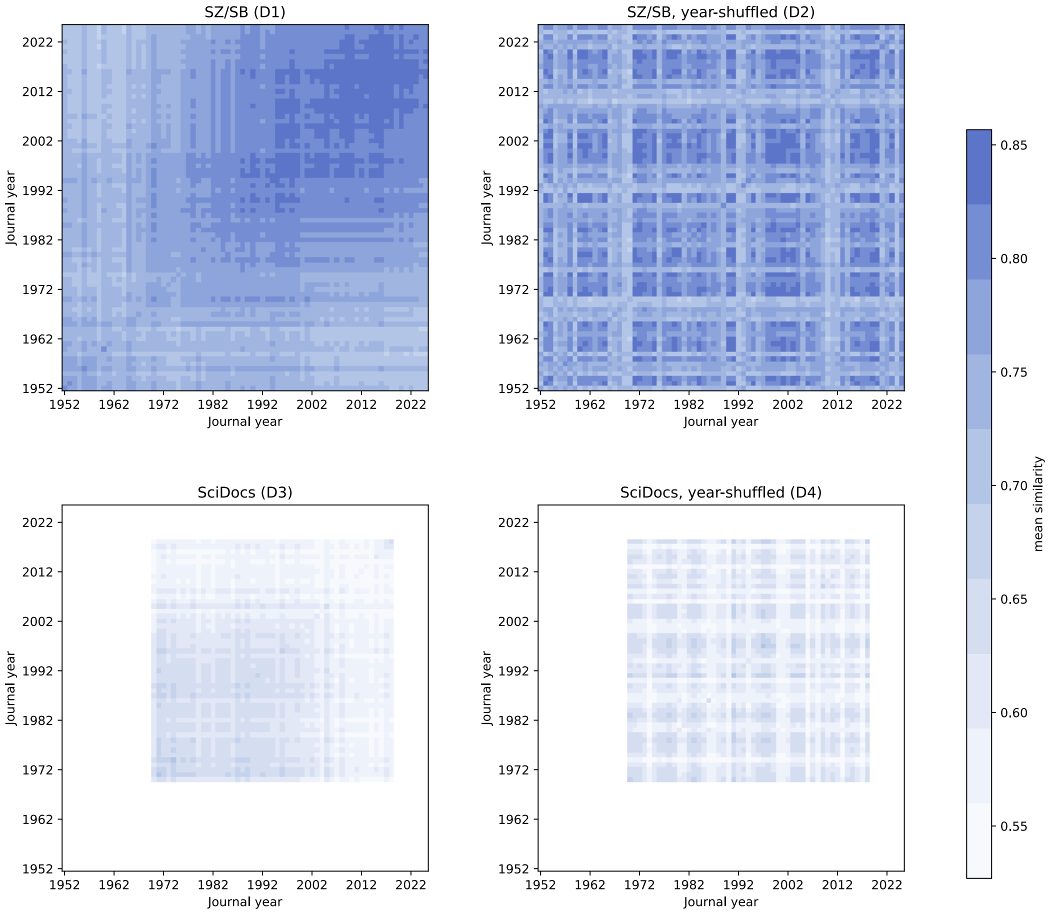


**Supplementary Figure S1.** Mean similarity among research articles between different publication years for the *Systematic Zoology* and *Systematic Biology* dataset with correct (*D1*) and shuffled (*D2*) publication years (1952-2024), and for the subsampled SciDocs dataset with correct (*D3*) and shuffled (*D4*) publication years (1970-2019). Similarity scores may range from 0 to 1. Note, the results for *D1* are identical to those presented in Figure 1 of the main text, except the color scale was altered to aid comparison, given the lower similarity scores generated by *D3* and *D4*. See Supplement text for details.

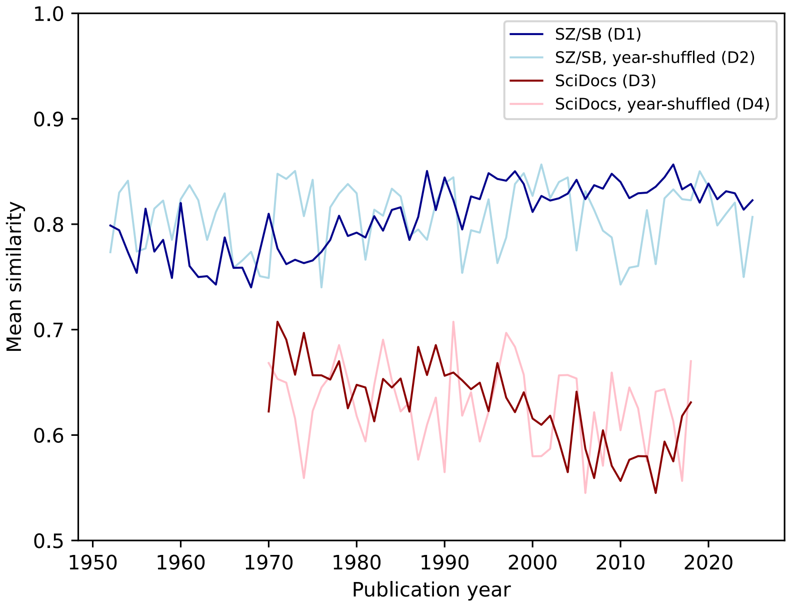


**Supplementary Figure S2.** Mean similarity among research articles within each publication year for the *Systematic Zoology* and *Systematic Biology* dataset with correct (*D1*) and shuffled (*D2*) publication years (1952-2024)*,* and for the subsampled SciDocs dataset with correct (*D3*) and shuffled (*D4*) publication years (1970-2019). Similarity scores may range from 0 to 1. The plotted similarity scores correspond to the diagonal cells of the heatmaps presented in Supplementary Figure S1. See Supplement text for details.

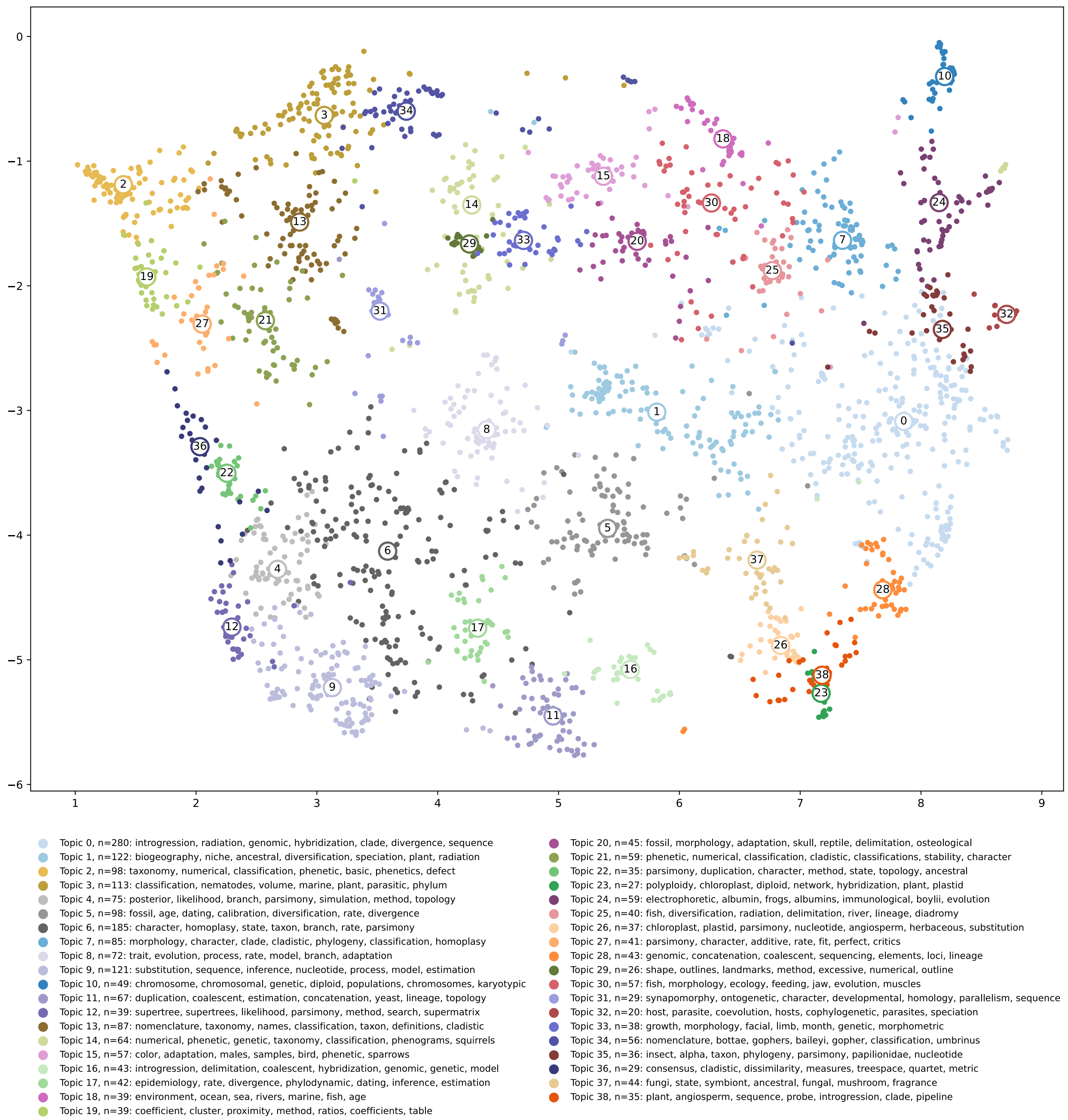


**Supplementary Figure S3.** *Systematic Zoology/Biology* research articles and topics. BERTopic associated 2,531 SPECTER-embedded research articles (points) with 39 inferred topics (colors), represented here in two dimensions using UMAP. See Supplement text for details.

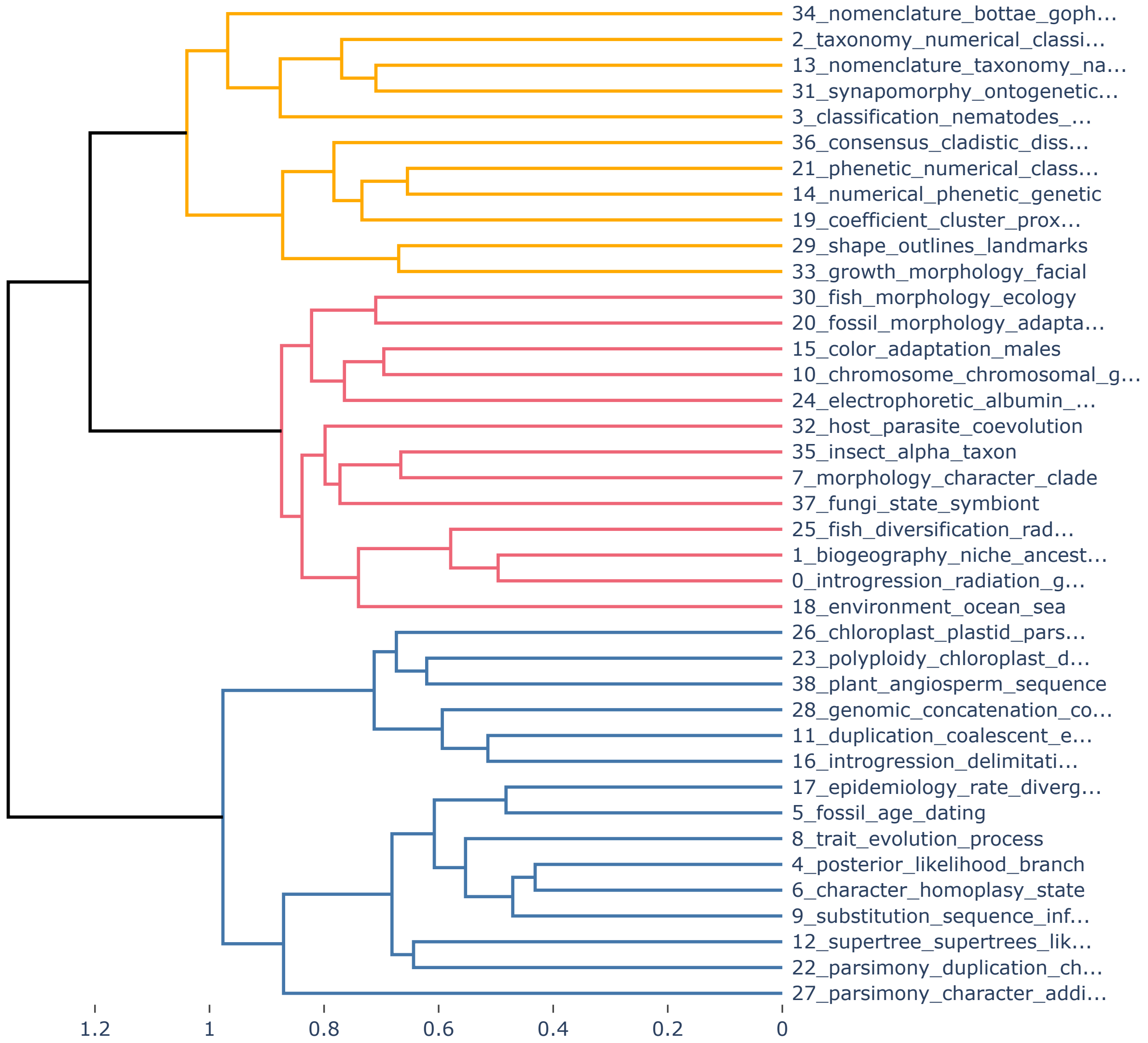


**Supplementary Figure S4.** Hierarchy of *Systematic Zoology/Biology* research article topics inferred by BERTopic. Cluster 1 (orange, top) generally contains topics associated with classification, nomenclature, and phenetics. Cluster 2 (red, middle) generally contains topics associated with organismal research ("clade biology"). Cluster 3 (blue, bottom) generally contains topics associated with models, methods, and phylogenomic approaches ("phylogenetic biology"). These main clusters are also used in Figure 1 of the main text and Supplementary Figures S6 and S7. Distances are measured as (1 – cosine similarity). See Supplement text for details.

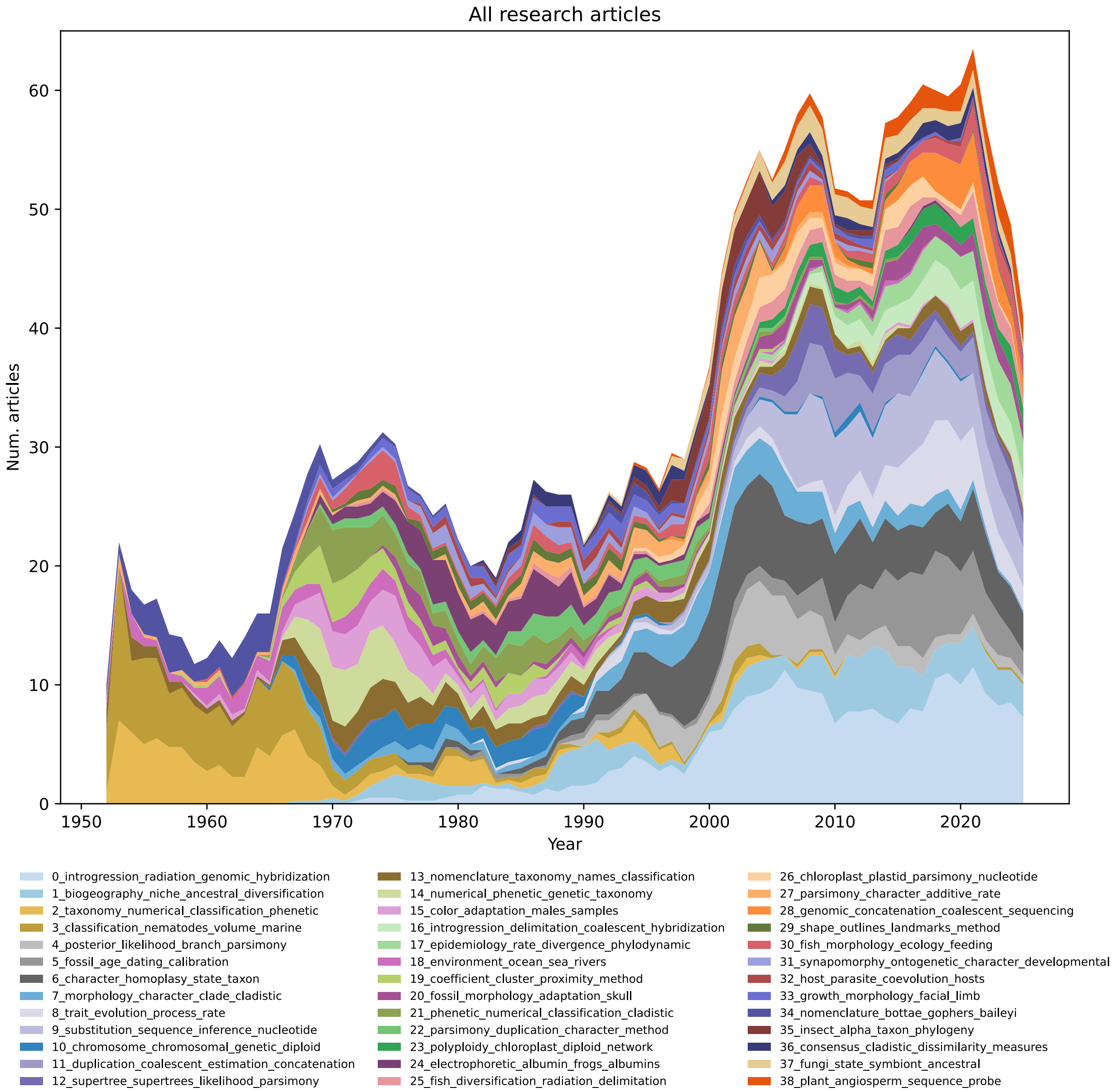


**Supplementary Figure S5.** Numbers of research articles per topic published in *Systematic Biology* during 1952 - 2025. BERTopic topic prediction divided articles into 39 topics (colors). Article counts were smoothed with a 5-year moving average to improve readability. See main text and supplementary methods for full details. Supplementary Figure S3 displays all topics. Figure 1 in the main text shows the trajectories of topics for each cluster represented in Supplementary Figure S4. See Supplement text for details.

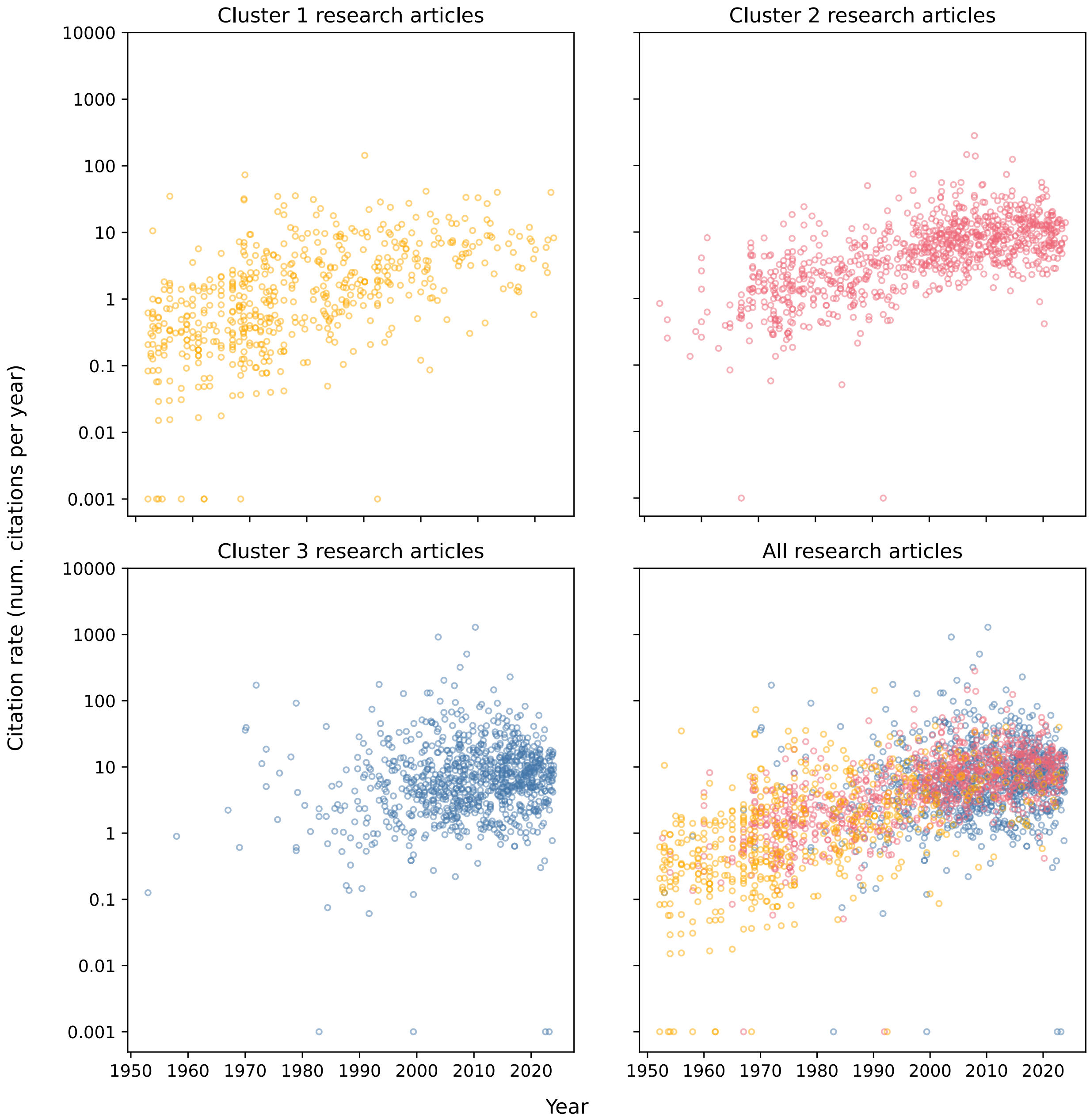


**Supplementary Figure S6.** Citation rates through time for research article topic-clusters. Cluster 1 (gold, upper left) generally contains topics associated with classification, nomenclature, and phenetics. Cluster 2 (red, upper right) generally contains topics associated with organismal research (“clade biology”). Cluster 3 (blue, bottom left) generally contains topics associated with models, methods, and phylogenomic approaches (“phylogenetic biology”). All articles are displayed in the lower right. Articles with citation rates lower than 0.001 were rounded up to that value. See Supplement text for details.

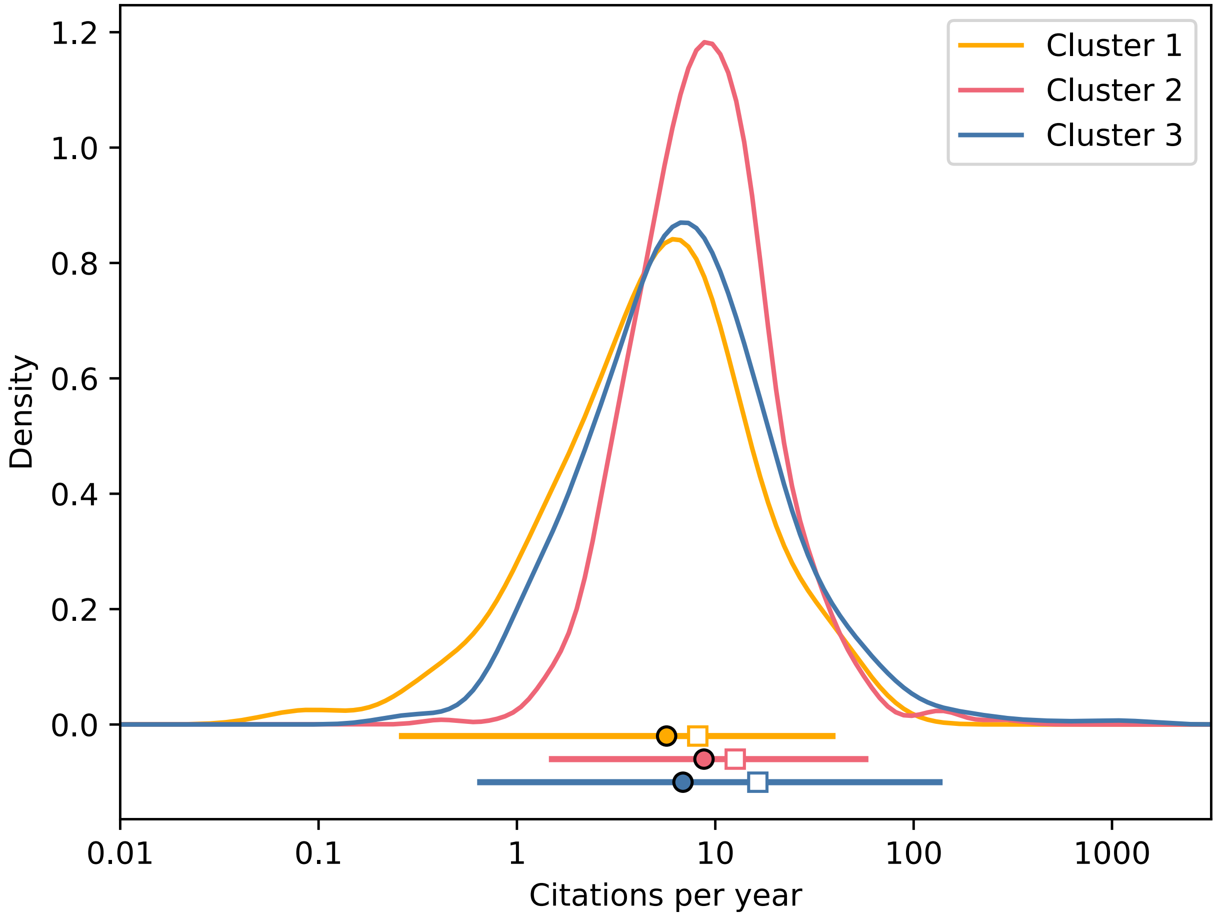


**Supplementary Figure S7.** Citation rates for research articles published during 2000-2024 per topic-cluster. Citation rate is computed as total number of citations divided by article age. Cluster 1 (gold, top) generally contains topics associated with classification, nomenclature, and phenetics. Cluster 2 (red, middle) generally contains topics associated with organismal research (“clade biology”). Cluster 3 (blue, bottom) generally contains topics associated with models, methods, and phylogenomic approaches (“phylogenetic biology”). Citation rate densities are accompanied by means (open squares), medians (filled circles), and 1% and 99% quantiles (horizontal bars). Articles with citation rates less than 0.01 citations/year not shown. See Supplement text for details.

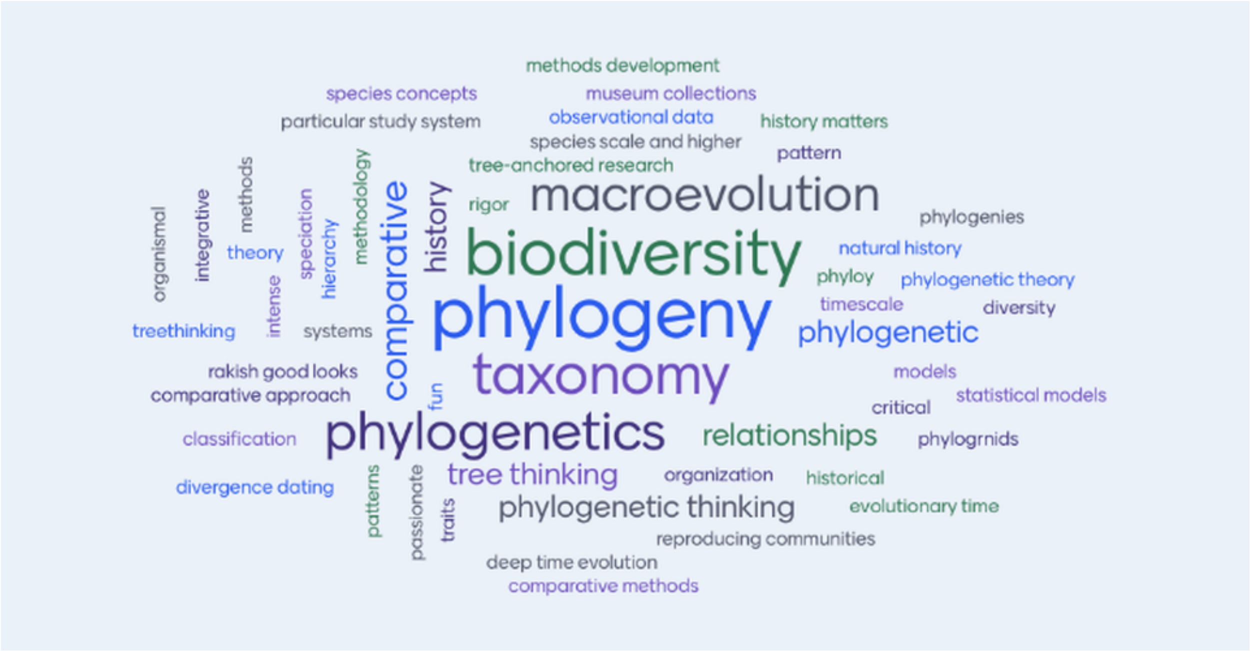


**Supplementary Figure S8.** Word cloud from the Society of Systematic Biologists home page that is referenced in the main text. The original caption for the image reads: "*Word Cloud from 2024 Members Meeting when participants were asked, 'What attribute do you believe distinguishes systematic biologists from other evolutionary biologists?'*". Image copied October 19, 2025.
